# Using multiple measurements of tissue to estimate subject- and cell-type-specific gene expression

**DOI:** 10.1101/379099

**Authors:** Jiebiao Wang, Bernie Devlin, Kathryn Roeder

## Abstract

**Motivation:** Patterns of gene expression, quantified at the level of tissue or cells, can inform on etiology of disease. There are now rich resources for tissue-level (bulk) gene expression data, which have been collected from thousands of subjects, and resources involving single-cell RNA-sequencing (scRNA-seq) data are expanding rapidly. The latter yields cell type information, although the data can be noisy and typically are derived from a small number of subjects.

**Results:** Complementing these approaches, we develop a method to estimate subject- and cell-type-specific (CTS) gene expression from tissue using an empirical Bayes method that borrows information across multiple measurements of the same tissue per subject (e.g., multiple regions of the brain). Analyzing expression data from multiple brain regions from the Genotype-Tissue Expression project (GTEx) reveals CTS expression, which then permits downstream analyses, such as identification of CTS expression Quantitative Trait Loci (eQTL).

**Availability and implementation:** We implement this method as an R package MIND, hosted on https://github.com/randel/MIND.

## 1 Introduction

Altered gene expression is one mechanism by which genetic variation confers risk for complex disease. Thus, many studies have quantified bulk gene expression from tissue, thereby assessing expression averaged over the individual cells comprising the tissue. Recently, using single-cell RNA sequencing (scRNA-seq) (Darmanis *et al.*, 2015; Zeisel *et al.*, 2015; Habib *et al.*, 2017), studies have quantified gene expression at the level of cells and cell types; such data could be especially informative for brain tissue, which harbors myriad cell types whose functions are not fully resolved. Drawbacks to scRNA-seq data include its noisy nature and the challenge of characterizing such cells from many subjects, which limits its potential for genetic analyses. Alternatively, there are established resources, such as GTEx(GTEx Consortium, 2017) and BrainSpan(Kang *et al.*, 2011), among others, that have collected bulk transcriptome data from many subjects and multiple brain regions. Although bulk transcriptomes represent an amalgamation of different cell types, which occur in various proportions in the sampled tissues, there is cell-type-specific information encoded in these transcriptomes, as recent studies demonstrate (Kelley *et al.*, 2018; Wang *et al.*, 2018). Here we present a method, MIND for Multi-measure INdividual Deconvolution (Fig. 1), to exploit such resources to learn about subject- and cell-type-specific (CTS) gene expression. For each subject and gene, MIND’s CTS estimate represents the average expression of the gene for fundamental cell types, such as neurons, astrocytes, microglia, oligodendrocytes and endothelial cells in brain.

**Fig 1.**
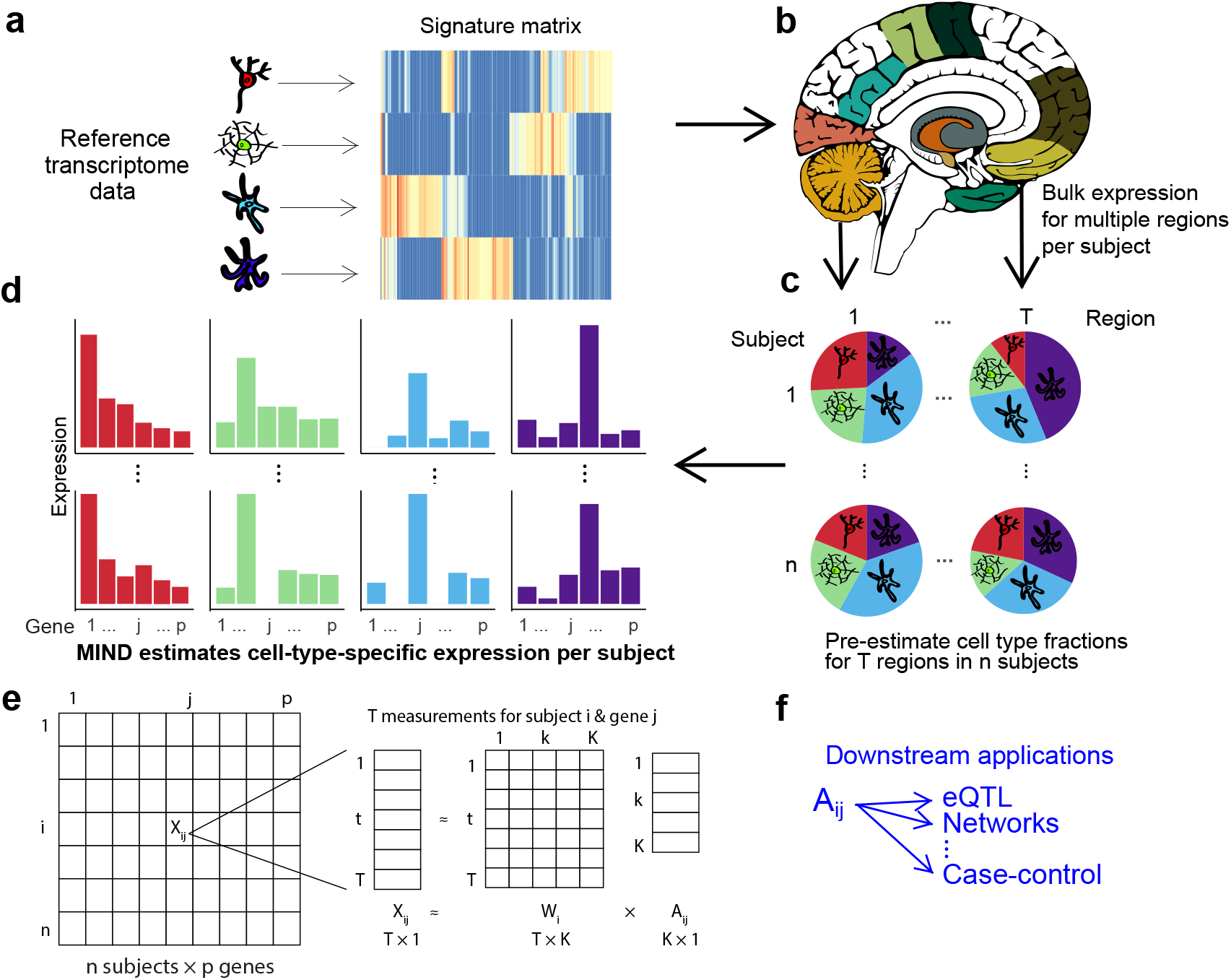
Flow diagram for the MIND algorithm. (a) For a set of relevant cell types, select cell type marker genes and build a signature matrix using reference samples. (b) Multiple transcriptomes are measured from each subject; here, one transcriptome for each of multiple brain regions. (c) Using an existing deconvolution method, e.g., non-negative least squares, estimate the cell type fractions for each brain region and subject. Here we depict *K* = 4 cell types for which their fractions will be estimated per brain region. (d) With results from (b) and (c), MIND estimates cell-type-specific (CTS) expression for each of *p* genes for each subject and cell type. Colors map to the cell types in (c) and (d) and we depict two of n subjects, 1 and *n*. (e) Matrix representation of key data elements of the MIND algorithm: for each of *T* brain regions for subject *i*, expression of *p* genes from the transcriptome is measured, ***X**_ij_*; and the key outputs are the subject level CTS gene expression (***A**_i_*) and the subject and measurement level cell type fractions (***W**_i_*). (f) Examples of downstream applications for MIND.

MIND exploits two key ideas for obtaining CTS gene expression from tissue. (1) Because a tissue sample’s bulk transcriptome is a convolution of gene expression from cells belonging to various cell types, deconvolution methods (Abbas *et al.*, 2009; Shen-Orr *et al.*, 2010; Newman *et al.*, 2015; Wang *et al.*, 2019) can estimate the fraction of each cell type within this tissue. Methods typically deconvolve each tissue sample per subject and require prior information, specifically sets of genes that are expressed in certain cell types (marker genes), the collection of which we call the signature matrix. Any good deconvolution method will suffice for MIND, such as CIBERSORT (Newman *et al.*, 2015) or the non-negative least squares approach adopted by PsychENCODE (Wang *et al.*, 2018). (2) Multiple transcriptomes from the same subject, but different brain regions, share common cell types. MIND uses empirical Bayes techniques to exploit this commonality, together with the estimated cell type fractions from (1), to estimate CTS gene expression. Note that the idea behind (2) is unique to MIND and it permits subject and CTS expression estimates from tissue level transcriptomes. Using MIND, we analyze data from GTEx brain tissue to obtain CTS gene expression.

## 2 Methods

### 2.1 The MIND algorithm

For a single measure (*t*) from subject *i*, let *X_ijt_* be the observed expression of gene *j*. When the tissue consists of *K* cell types, typically the goal of gene expression deconvolution is to find ***W**_it_*, the *K* cell type fractions for subject *i* in measure *t*, such that

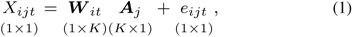

where ***A**_j_* is the cell type gene expression and *e_ijt_* is the error term. When reference samples are available, such as purified cells or scRNA-seq data, the signature matrix can be estimated for the marker genes by differential expression analysis of cell types from the reference samples. Plugging in ***A_j_***, deconvolution becomes a standard regression problem and ***W**_it_* can be estimated directly.

We extend the single-measure deconvolution in equation (1) by borrowing information across multiple measurements, *t* = 1,…, *T_i_* from the same tissue for subject *i* to estimate subject-specific and CTS gene expression (*T_i_* can vary by subject.)

**Step 1** of the MIND algorithm is to estimate cell type fractions for subject *i* and measure *t*, ***W**_it_*, for *t* = 1,…, *T_i_*. Combining estimated information across measures yields ***W**_i_*, a *T_i_* × *K* matrix, of cell type fractions. **Step 2**, treating ***W**_i_* as known, we reverse the problem from single-measure deconvolution, estimating instead CTS gene expression. For gene *j* in subject *i*, the observed gene expression ***X**_j_* is a *T_i_* × 1 vector that represents *T_i_* quantified measurements (Fig. 1e), rather than a scalar as in equation (1). We model ***X**_ij_* as a product of cell type fraction (***W**_i_*) and CTS expression (***A**_ij_*),

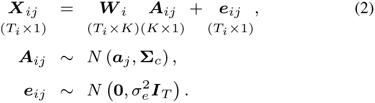

where *e_ij_* is the error term that captures the unexplained random noise.

To ensure robustness, we assume that the CTS expression (***A**_j_*) is randomly distributed as ***A**_ij_* ~ *N* (***a**_j_*, **Σ**_*c*_), with mean ***a**_j_* (*K* × 1) that constitutes the profile matrix and covariance matrix **Σ**_*c*_ (*K* × *K*) for *K* cell types. In contrast to single-measure deconvolution, we assume

1. cell type fraction (***W**_i_*) is subject- and measure-specific;
2. CTS expression (***A**_ij_*) is subject-specific but constant across measures.

Estimation is performed across all subjects and genes simultaneously. We estimate the parameters (***a**_j_*, **Σ**_*c*_) through maximum likelihood via a computationally efficient EM (Expectation-Maximization) algorithm (see Supplementary Note). CTS expression (***A**_ij_*) is estimated using an empirical Bayes procedure. To achieve reliable results, the number of cell types (*K*) to be estimated is limited by the number of measures (e.g., brain regions) per subject, whereas all genes in the genome can be efficiently deconvolved together. MIND ignores gene-gene correlation in the prior distribution of CTS expression to achieve efficient computation, deconvolving for the whole genome in several minutes (Supplementary Table 1). Gene-gene correlation can be inferred from estimates of CTS gene expression.

### 2.2 Data resources, preliminary analyses, simulations

To explore the performance of MIND, we estimated subject and CTS gene expression from GTEx bulk gene expression data from human brain (GTEx Consortium, 2017). GTEx is an ongoing project that collects both gene expression data from multiple tissue types, including brain, and genotype data from blood for hundreds of post-mortem adult donors. Here we focused on 1671 brain tissue samples from 254 donors and 13 brain regions in the GTEx V7 data (GTEx Consortium, 2017). We analyzed the read count data for all genes detected in brain, normalized as count per million (CPM) and transformed as log_2_(*X* + 1) prior to analysis. Unless otherwise noted, all expression count data analyzed herein were log-transformed in this way. (See Supplementary Note for discussion and analyses of various transformations of the data.)

Samples of brain tissue from different brain regions share common cell types and thus are suitable for analysis by MIND. To ensure reliable estimates, we removed GTEx subjects with less than 9 brain regions sampled, resulting in data from 105 subjects for analysis. Among these subjects, 95 also had genotype data for identifying CTS eQTLs. These eQTLs were estimated using MatrixEQTL (Shabalin, 2012), controlling for ancestry, gender, and genotype platform and were compared to eQTLs from GTEx bulk data, specifically region-specific eQTLs downloaded from the GTEx portal.

To build a signature matrix for the GTEx brain data, we began by analyzing the adult scRNA-seq data from Darmanis *et al.* (2015), clustering characterized human brain cells into seven cell types (astrocyte, endothelial, microglia, oligodendrocyte (Oligo), oligodendrocyte precursor cell (OPC), inhibitory and excitatory neurons) and selecting the top 50 marker genes for each cell type using SC3 (Kiselev *et al.*, 2017). We also collected markers for astrocyte, microglial, and endothelial cells from the PsychENCODE Consortium (Wang *et al.*, 2018) and microglia markers from Olah et *al.* (2018). The signature matrix (Supplementary Table 2) was constructed by averaging the expression of each marker gene in the cells of the same type. Following the PsychENCODE Consortium (Wang *et al.*, 2018), we estimated the cell type fractions for GTEx subject-by-brain region using non-negative least squares (Lawson and Hanson, 1995) (Supplementary Table 3). Because the estimated fractions of OPCs in brain regions were always close to zero, OPCs were dropped from further analyses. Given estimated cell type fractions, MIND then estimates subject and CTS gene expression from GTEx bulk expression data.

MIND’s estimates were validated empirically and by simulation. For empirical validation, we used data from Habib et al. (2017), who quantified single-nucleus RNA-seq data from seven brain tissue samples from five GTEx donors. Because the authors classified the cells into cell types, we could average their read count data for cells of each type to obtain CTS expression on a scale similar to that produced by MIND. Because only a few hundred cells were characterized for one of the five subjects and these data cannot provide accurate CTS expression, data from this subject were excluded. For a fair comparison, we converted the read counts to CPM and then compared the directly measured subject-specific and CTS expression to MIND’s estimated quantities from bulk transcriptomes (in CPM) from the same subjects.

To evaluate MIND via simulations, we generated gene expression using two approaches. To enhance the realism, for each approach we used cell type fractions (***W**_i_*) estimated from GTEx subjects, as described above, for 6 cell types (astrocyte, oligodendrocyte, microglia, endothelial cell, inhibitory, and excitatory neurons). As a measure of performance, we computed the correlation between MIND’s and the true CTS expression. To evaluate the parameter estimates, we calculated the average correlation and variance estimate from 100 replications. In the first simulation approach, using equation (2), we simulated 105 subjects having 9-13 measurements as in GTEx brain data. We utilized the profile matrix (***a**_j_*) built from Darmanis *et al.* (2015) for two sets of genes: (i) markers gene, and (ii) all genes. We systematically varied the values of the true variance parameters, 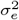 and **Σ**_*c*_, which denote the error variance and the covariance of CTS expression. Here we let **Σ**_*c*_ have equal variance 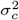 and equal covariance 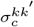 across cell types, where *k* and *k′* denote cell types. For the second simulation approach, single-cell data from 4 GTEx subjects, described above, guided data production for 100 subjects and 31,496 genes. There were two settings in the second simulation approach, (1) we varied the error variance 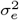 and (2) we added region-specific variation to ***A**_ij_* (CTS expression per subject), letting the variation follow a normal distribution with zero mean and variance equal to the simulated error variance 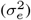. Finally we fixed 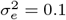 (the error variance that we observed in deconvolving GTEx brain data) and varied the number of measures from 1 to 13.

To partition variation in gene expression by cell type and brain region and thereby check MIND assumptions, we used the NeuroExpresso database (Mancarci *et al.*, 2017), which holds gene expression data for purified-cell samples from multiple mouse brains and regions. This resource holds normalized data of purified cells on expression of 11,546 genes. We restricted our analysis to four cell types with largest sample size, namely astrocytes, oligodendrocytes, and inhibitory/GABAergic and excitatory/pyramidal neurons.

## 3 Results

### 3.1 Validating model assumptions

MIND models cell type fraction as subject- and region-specific. It is natural to assume CTS expression is subject-specific, which allows for differences among subjects due to age, phenotype, genotype and other measured variables and thereby permits downstream analyses not formerly possible (Fig. 1f). MIND also assumes CTS expression is similar across brain regions of the same subject, thereby avoiding overfitting the data. For this assumption to hold, cells from the same cell type, but from different brain regions, should show similar patterns of gene expression; whereas cells of different cell types from the same region should show distinct expression profiles. This was the observed pattern in the NeuroExpresso database of purified brain cells from multiple brain regions (Fig. 2a). Fitting a mixed-effects model for each gene and decomposing the variance into that explained by cell types versus brain regions, as well as studies and error, cell types account for a larger amount of the variance than region, 25% versus 12%.

**Fig 2.**
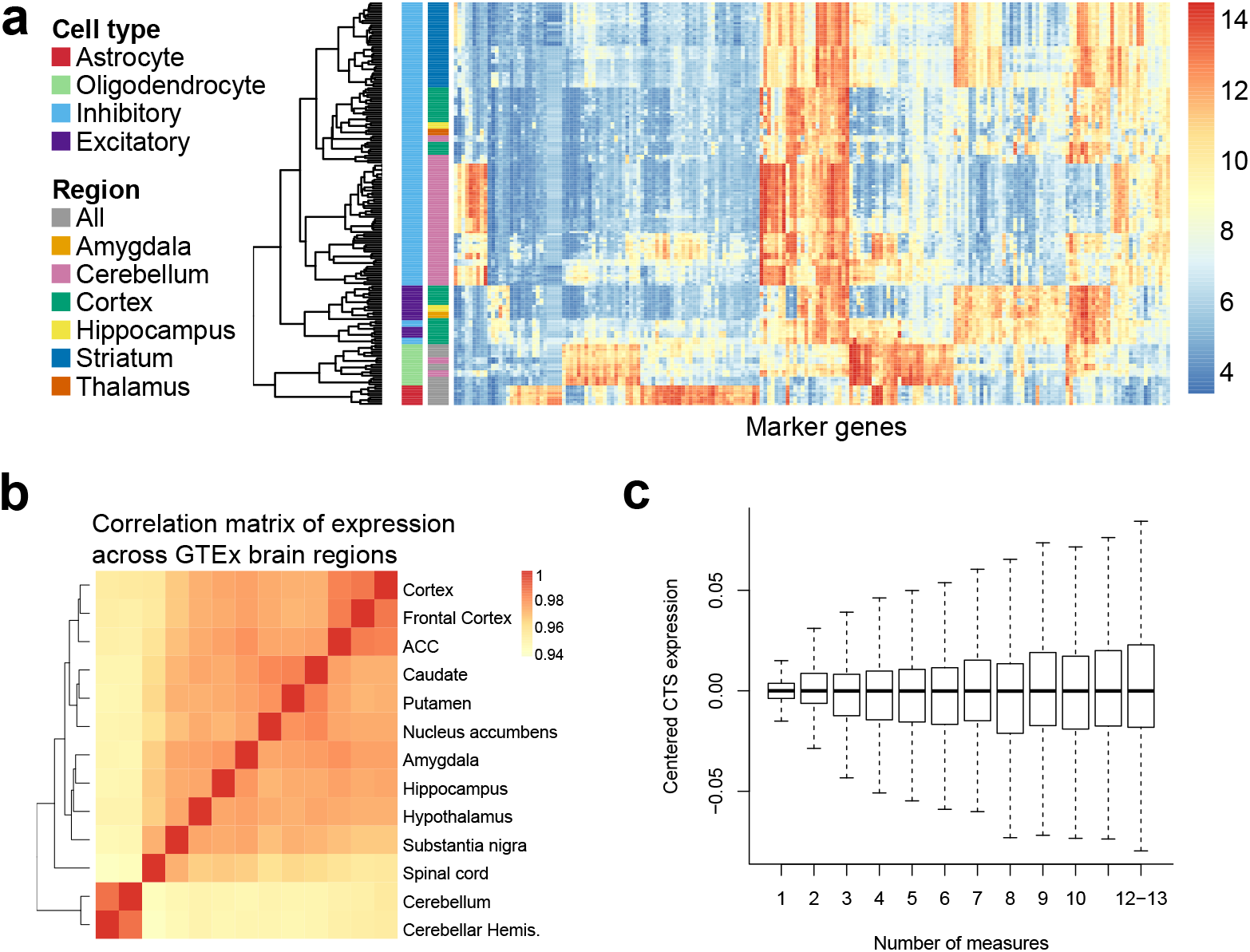
Validation of the assumptions of MIND. (a) Heatmap of expression of cell type marker genes in the NeuroExpresso database of purified-cell samples. Columns denote 192 marker genes selected by CIBERSORT (Newman *et al.*, 2015) from NeuroExpresso. Rows represent purified-cell samples. Purified-cell samples are clustered, then annotated by cell type and brain region (labels on left, scale of expression on right). (b) Correlation matrix of gene expression (heatmap) for brain regions from GTEx samples. (ACC: anterior cingulate cortex; hemis.: hemisphere.) (c) Boxplots of centered CTS expression as a function of the number of measures per subject in GTEx brain data. The number of subjects with all 13 measures is small and thus those subjects are combined with subjects having 12 measures. To obtain centered CTS expression, for a given number of measures, the estimated expression per gene and cell-type was first centered, then these estimates were pooled for display.

Next, examination of the correlation of gene expression over regions for the GTEx brain data shows that bulk gene expression is highly correlated over all regions, with cerebellum and spinal cord showing slightly lower correlation (Fig. 2b). Reversing the role of region and subject in MIND, to estimate CTS expression for every region, shows that the estimated expression is quite similar across regions as illustrated by marker genes (Supplementary Fig. 1), with the strongest deviation observed for cerebellum. Fitting a mixed-effects model for each gene and decomposing the variance into that explained by cell types and brain regions, the variance explained by cell types (41%) is substantially larger than that for regions (1%).

It is reasonable to ask if MIND requires repeated measures of gene expression in the same or similar tissue. Results using the deconvolved GTEx brain data show that for subjects with fewer measures, the deconvolved CTS expression has less variability, on average (Fig. 2c). This implies that the model typically imputes similar expression for each cell type when the number of measurements is small and it lacks strong information to the contrary. Thus the number of measurements provides an indicator of the reliability of the deconvolved expression.

### 3.2 Validating model estimates

We evaluate the performance of MIND for various scenarios, including a variety of simulations and analysis of the GTEx brain tissue data. Importantly, GTEx (Habib *et al.*, 2017) produced scRNA-seq data from the prefrontal cortex (3 subjects) and hippocampus (4 subjects). For these same subjects, bulk transcriptomes were also characterized (GTEx Consortium, 2017). From the scRNA-seq data, we can calculate CTS expression by averaging over cells of each cell type for each subject. Then, existence of both bulk and scRNA-seq data enables a direct comparison of MIND’s performance and reveals highly concordant estimates for most cell types and donors (Fig. 3a). As expected, focusing on marker genes, we find that MIND’s CTS expression has the highest correlation with the measured expression of the same cell type (Supplementary Table 4). If MIND’s estimates are accurate, bulk gene expression should be a convolution of its estimated CTS gene expression and the estimated cell type fraction for the tissue sample. Using MIND’s estimates to predict region level expression for each subject shows excellent correspondence between predicted versus measured bulk gene expression (Supplementary Fig. 2).

**Fig 3.**
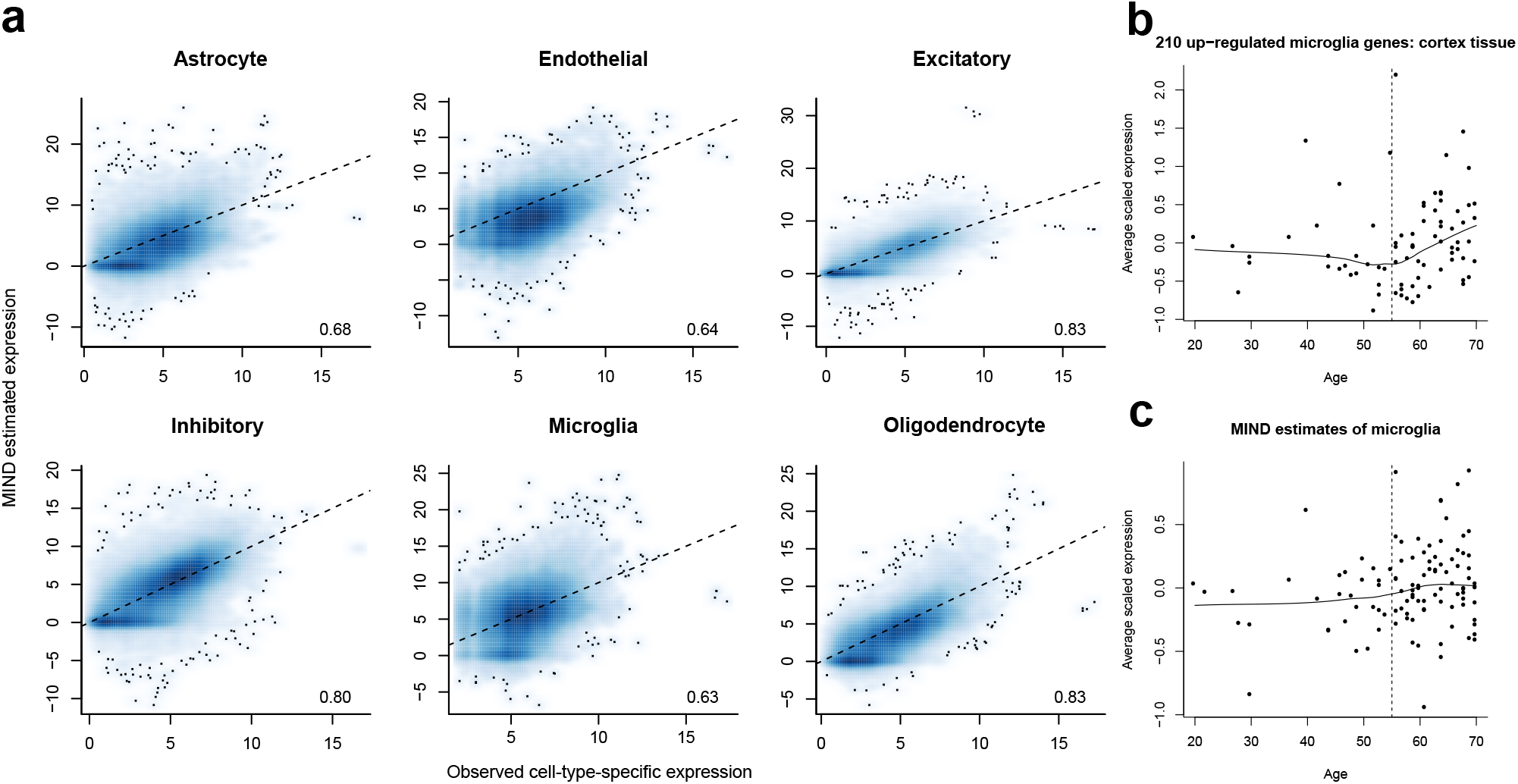
Validation of the estimates of MIND. (a) Direct quantification of average gene expression from single cells (observed) (Habib *et al.*, 2017) from GTEx brain samples of the same subjects as the CTS expression estimated by MIND. Shown are scatter plots represented as a smoothed two-dimensional color density. For each panel, correlation for all genes is given. For presentation, only genes with positive observed expression are shown. On average, there are 17,223 out of 31,496 genes that have positive observed CTS expression. Dotted line at *y* = *x*. (b) Average scaled expression of microglia marker genes in GTEx cortex tissue and matching the pattern observed by Olah et al. (2018) from different subjects. (c) MIND derived average scaled expression of the same microglia marker genes analyzed in (b) and showing the same pattern of increased expression with age. Expression values in (b) and (c) have been centered.

Recent research identifies microglia as a cell type involved in risk for Alzheimer disease. In this regard, Olah *et al.* (2018) identified a set of genes that mark microglia in the aging human brain and showed expression increased with age. We replicate the same pattern in GTEx brains (Fig. 3b) and, importantly, we see the same trend in MIND’s estimate of the expression of these genes in microglia (Fig. 3c).

The design of simulation studies to evaluate MIND’s performance was described in Methods. In Supplementary Figure 3 and Table 5, we provide detailed results of these studies. In brief, MIND provided consistently high correlation between estimated and true CTS gene expression for all cell types and was robust to increasing noise (Supplementary Fig. 3a). This conclusion held even when we simulated bulk data with region-specific CTS expression (Supplementary Fig. 3b) and MIND’s performance improved with the number of regions assessed (Supplementary Fig. 3c). When six or more regions were assessed, the correlation between the estimated and true CTS expression for six cell types reached 0.8 on average when the error variance was set equal to that observed from deconvolved GTEx brain data. MIND provided high quality estimates when the cell type fraction is ≥ 0.05, but it cannot recover accurate estimates of CTS expression for more rare cell types (Supplementary Fig. 3d). Moreover, MIND’s parameter estimates were approximately unbiased (Supplementary Table 5) when the number of measures was large. A least-squares approach did not perform as well as MIND (Supplementary Fig. 3), highlighting the advantages of considering correlations between measures and assuming random CTS expression in MIND, an assumption that is particularly valuable when the number of measures is small, which is usually the case in practice.

### 3.3 Further results from GTEx brain tissue

Results for cell type fractions (Fig. 4a) were consistent with previous findings and what is known about the brain: (i) related brain regions have similar cell type composition, for example, the three basal ganglia structures, two cerebellum samples, and three cortical samples (note that the two cerebellum samples and two frontal cortical samples are technical replicates); and (ii) spinal cord (cervical c-1) is estimated to consist of 35% oligodendrocytes, which agrees with the prominence of white matter tracts present at c-1 and glial cells in white matter. See Supplementary Note for analyses and discussion of the interrelationship of estimated cell fractions, cell size and gene expression (Supplementary Fig. 4).

**Fig 4.**
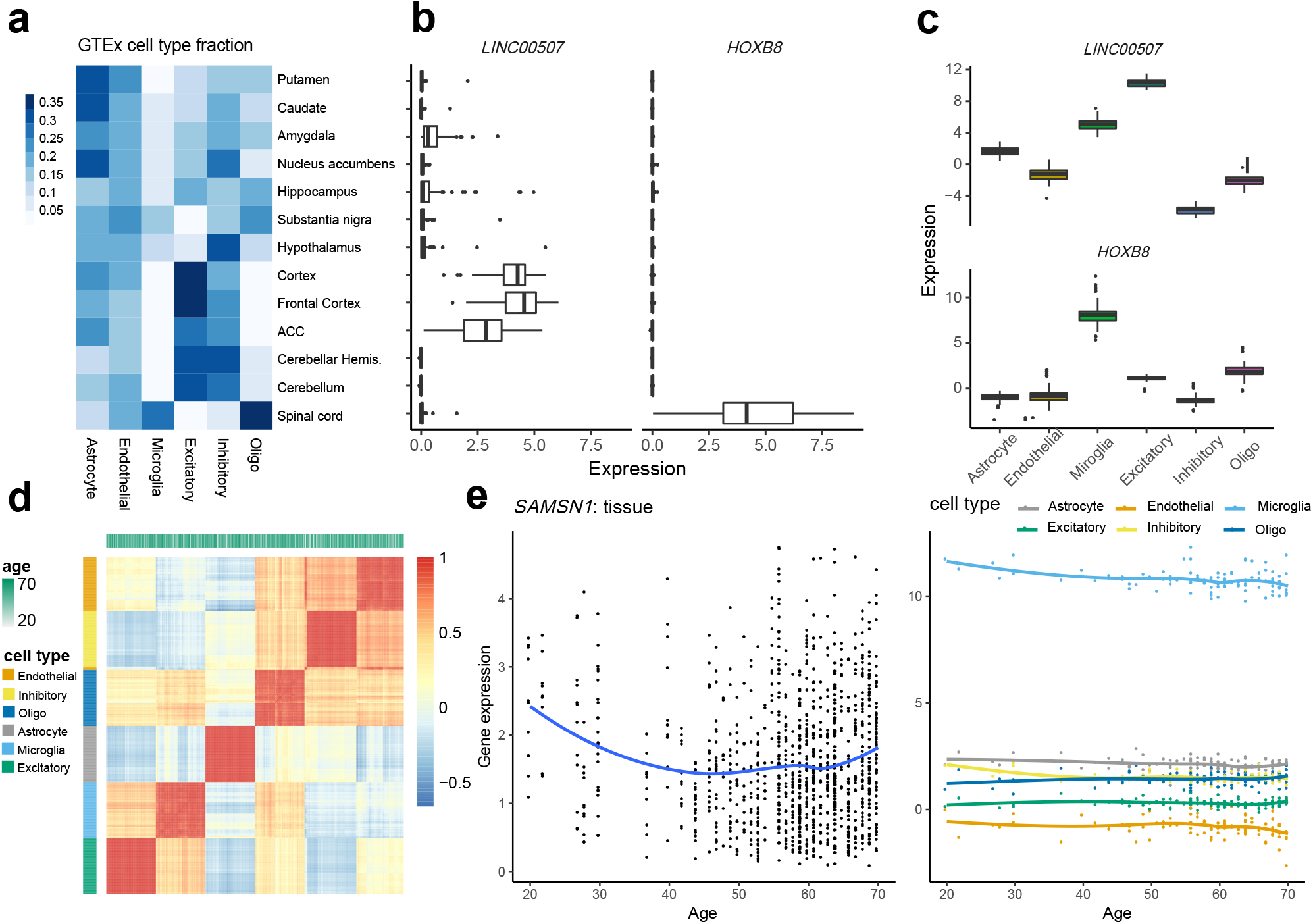
Analyses of CTS gene expression of the GTEx brain data. (a) Estimated cell type fractions in each GTEx brain region, averaged over subjects. Putamen, caudate, and nucleus accumbens are the three basal ganglia structures. (b-c) For two transcripts selected for differential expression in cortex versus spinal cord, (b) boxplots of tissue-level expression across brain regions and (c) CTS expression estimated by MIND from tissue-level expression across brain regions. (d) The heatmap and clustering of estimated CTS expression from MIND by cell type and age. Here we visualize a 6*n* × 6*n* correlation matrix for the 6 cell types and *n* = 105 subjects, based on the expression of 124 genes that have the largest variability across brain regions. (e) Age trends for expression of gene SAMSN1 in tissue and its estimated CTS expression from MIND. SAMSN1 is a known marker gene for microglia and is down-regulated in aged microglia (Olah *et al.*, 2018).

We next examine the estimated CTS expression values, by subject, to determine if the estimates conform to expected patterns. It is reasonable to predict that RNA showing specificity for certain brain regions would also show specificity to a cell type prominent in that region. This is indeed the case. For example, consider *LINC00507* and *HOXB8*, the former is highly expressed in cortical brain tissue, and the latter in spinal cord (Fig. 4b). By contrasting the region-level expression for these genes with their estimated CTS expression (Fig. 4c), we find that *LINC00507* tends to be expressed solely in excitatory neurons (Aevermann *et al.*, 2018), while *HOXB8* is expressed largely in microglia (Frick and Pittenger, 2016). A priori, and based on recent findings (Soreq *et al.*, 2017), we would also expect cell type to be a strong predictor of gene co-expression. Moreover, because GTEx subjects were all adults at death, but not elderly, recent findings (Soreq *et al.*, 2017) suggest that age would not be a strong predictor of gene co-expression. Thus, we asked if the estimated CTS expression clusters by cell type or by age of the subject using estimates from 124 genes with the largest variability in expression across brain regions. Based on these genes, we compute the correlation matrix for the 6*n* subject-cell-type configurations (6 cell types and *n* = 105 subjects). Hierarchical clustering of the entries in the correlation matrix reveals that cell-type is a strong predictor of co-expression, while age is not (Fig. 4d), consistent with MIND’s modeling assumptions.

CTS expression by age, however, reveals interesting patterns that are not always apparent at the tissue level. For example, for *SAMSN1*, expression decreases with age in tissue, whereas it is only expressed substantially in microglia and only shows a significant decreasing trend in microglia and inhibitory neurons (Fig. 4e). This agrees with findings in Olah *et al.* (2018) that *SAMSN1* is a marker for microglia and down-regulated in aged microglia cells. Overall, 21% of genes show age trends at the region level or cell-type level, with the false discovery rate (FDR) (Storey and Tibshirani, 2003) controlled at 0.05: 5% show age trends in at least one brain region and at least one cell type; 14% show age trends in at least one of 13 brain regions, but not in any cell type; and 2% show age trends in at least one of 6 cell types, but not in any brain region.

Because MIND yields subject-level and CTS gene expression, we can identify eQTLs for each cell type. To do so, CTS gene expression data were analyzed using MatrixEQTL (Shabalin, 2012), with FDR controlled at 0.05 for each cell type. We assessed that the p-values of eQTL mapping are well calibrated and enriched near gene transcriptional start site (Supplementary Fig. 5). We then compared the MIND-identified eQTLs with region-specific eQTLs identified by the GTEx project (GTEx Consortium, 2017). Notably, the rate at which eQTLs are both region-specific and CTS increases as the cell type becomes more prominent in the region (Fig. 5a). Moreover, when an eQTL was jointly identified in more brain cell types, it was more likely to be detected across a variety of tissues and especially across brain regions (McKenzie *et al.*, 2014) (Fig. 5b, Supplementary Fig. 6a). Interestingly, this does not hold for all cell types, because the eQTLs identified in oligodendrocyte and microglia were more distinct than those in the other four cell types (Supplementary Fig. 6b). We found that 48% of eQTLs that were identified in one or more brain cell types were not identified from any GTEx brain region, which suggests MIND’s results can identify novel eQTLs. Finally, some eQTLs were shared by all six cell types, while others are specific to certain cell types, especially for neuronal cells (Fig. 5c and Supplementary Fig. 6c), which implies that eQTL analysis based on MIND’s results can shed light on gene expression regulation within cell types. Additionally, those genes that had eQTLs in fewer cell types were more likely to be marker genes (Chi-squared-test of independence, p-value = 1.6 × 10^−28^).

**Fig 5.**
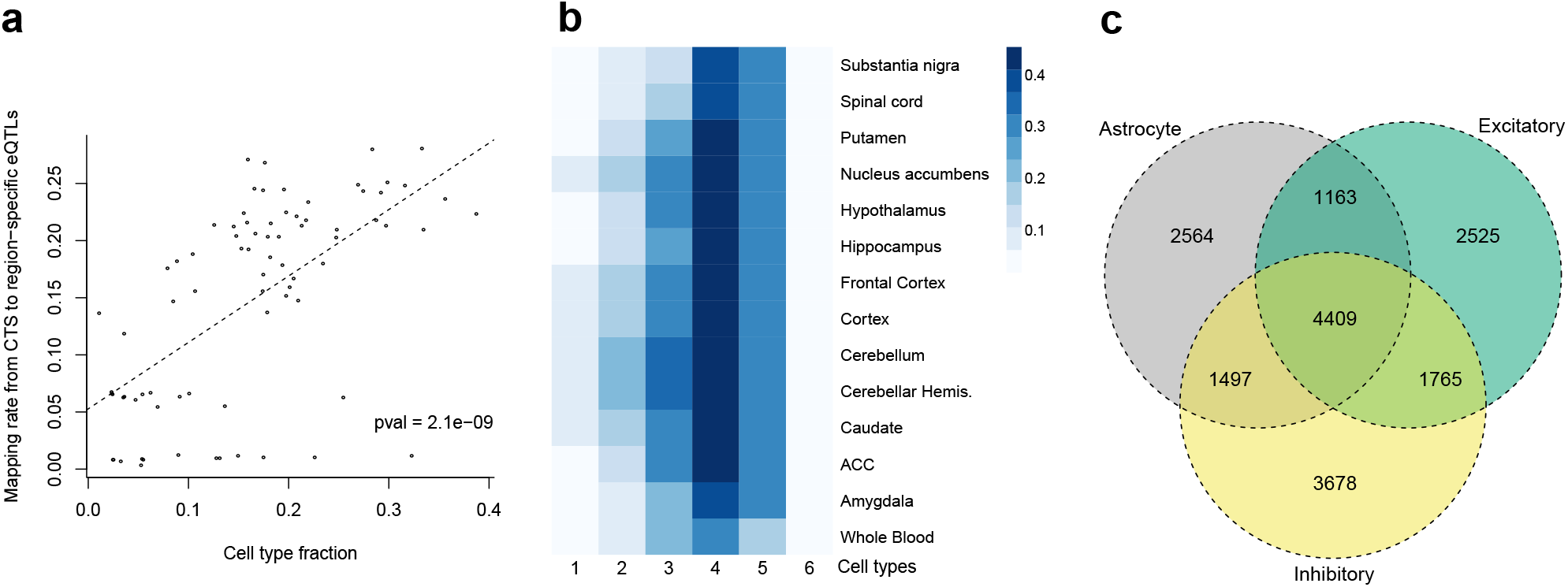
Expression quantitative trait loci (eQTL) discovered from tissue-level or CTS gene expression. (a) Scatter plot of eQTL mapping rate versus the estimated cell type fraction. The rate is for mapping eQTLs from CTS expression estimated by MIND to region-specific eQTLs identified by the GTEx consortium. It represents the probability of an eQTL detected in a cell type being identified in a brain region as well. Each point denotes a brain region and cell type. The dashed line depicts the fitted linear regression model and the p-value (pval) is for the t-test of the regression slope. (b) Rate of correspondence between eQTLs appearing in one to more cell types and those in each tissue type. For eQTLs that appear in one to six cell types, respectively, we calculate their probability of being identified in each brain region or whole blood. (c) Overlap among eGenes (genes with eQTLs) for three cell types of interest: astrocyte, inhibitory and excitatory neurons. The results for all six cell types are presented in Supplementary Fig. 6c.

## 4 Discussion

We develop an algorithm, MIND, to obtain gene expression by cell type and subject, even though gene expression is measured from tissue. There are notable advantages to the MIND algorithm. Because its estimates are CTS for each subject, they represent the cell-specific features inherent in the database for these subjects, such as the eQTLs from CTS expression. While we have concentrated our analyses on brain tissue, MIND is not specific to brain, any tissue could be appropriate, given these two conditions: there are a group of subjects for which transcriptomes have been assessed repeatedly; and the repeatedly sampled tissue, per subject, has cell types in common. For example, several other GTEx tissues meet these requirements, including artery and esophagus (GTEx Consortium, 2017). Other experimental settings fit these requirements too, such as organoids (Mariani *et al.*, 2015; Takasato *et al.*, 2015). It is also possible that one could substitute repeated measures per subject with repeated measures of genetically similar subjects, such as sibships for model organisms.

There are also limitations to the current version of MIND, which relies on reference samples to identify genes whose expression are largely specific to cell type, so-called marker genes. Identifying which reference samples are appropriate can be challenging. A different challenge is presented when there are a large number of cell types in the tissue. Reliably estimating expression by cell type and subject will require a large number of repeated measures per subject, something most resources do not have at this time. For this reason, we limit our analyses to major cell types. Furthermore, MIND is limited to estimating the average gene expression across cells of the same type within a subject, ignoring the diversity of expression within single cells.

Despite its limitations, MIND has the potential to increase the value of bulk transcriptome resources. By estimating individual and CTS gene expression, it can determine if an age trend for a gene’s expression is a property of a specific cell type or is a composite pattern arising from the various cell types comprising the tissue. Extending this observation, results from MIND can be used to produce CTS gene expression networks, which in turn can yield critical clues regarding the etiology of complex diseases. And, as documented here, it can map CTS variation in gene expression onto genetic variation, yielding CTS eQTLs, further enhancing the utility of bulk transcriptome resources to understand the origins of complex disease.

## Acknowledgements

We are grateful for the insightful comments from Michael Breen, Joseph Buxbaum, Lin Chen, Serkan Erdin, Dadi Gao, Lambertus Klei, Maria Jalbrzikowski, Silvia De Rubeis, Stephan Sanders, Michael Talkowski, and Haiyuan Yu, who read a previous version of the manuscript.

## References

Abbas, A. R., Wolslegel, K., Seshasayee, D., Modrusan, Z., and Clark, H. F. (2009). Deconvolution of blood microarray data identifies cellular activation patterns in systemic lupus erythematosus. PloS one, 4(7), e6098.

Aevermann, B. D., Novotny, M., Bakken, T., Miller, J. A., Diehl, A. D., Osumi-Sutherland, D., Lasken, R. S., Lein, E. S., and Scheuermann, R. H. (2018). Cell type discovery using single-cell transcriptomics: implications for ontological representation. Human molecular genetics, 27(R1), R40–R47.

Darmanis, S., Sloan, S. A., Zhang, Y., Enge, M., Caneda, C., Shuer, L. M., Hayden Gephart, M.G., Barres, B.A., and Quake, S. R. (2015). A survey of human brain transcriptome diversity at the single cell level. Proceedings of the National Academy of Sciences, 112(23), 7285–7290.

Frick, L. and Pittenger, C. (2016). Microglial dysregulation in ocd, tourette syndrome, and pandas. Journal of immunology research, 2016.

GTEx Consortium (2017). Genetic effects on gene expression across human tissues. Nature, 550(7675), 204.

Habib, N., Avraham-Davidi, I., Basu, A., Burks, T., Shekhar, K., Hofree, M., Choudhury, S. R., Aguet, F., Gelfand, E., Ardlie, K., et al. (2017). Massively parallel single-nucleus rna-seq with dronc-seq. NatureMethods, 14(10), 955–958.

Kang, H. J., Kawasawa, Y. I., Cheng, F., Zhu, Y., Xu, X., Li, M., Sousa, A. M., Pletikos, M., Meyer, K.A., Sedmak, G., et al. (2011). Spatio-temporal transcriptome of the human brain. Nature, 478(7370), 483.

Kelley, K. W., Nakao-Inoue, H., Molofsky, A. V., and Oldham, M. C. (2018). Variation among intact tissue samples reveals the core transcriptional features of human cns cell classes. Nature neuroscience, 21(9), 1171.

Kiselev, V. Y., Kirschner, K., Schaub, M. T., Andrews, T., Yiu, A., Chandra, T., Natarajan, K. N., Reik, W., Barahona, M., Green, A. R., et al. (2017). Sc3: consensus clustering of single-cell rna-seq data. Nature methods, 14(5), 483.

Lawson, C. L. and Hanson, R. J. (1995). Solving least squares problems, volume 15. Siam.

Mancarci, B. O., Toker, L., Tripathy, S. J., Li, B., Rocco, B., Sibille, E., and Pavlidis, P. (2017). Cross-Laboratory Analysis of Brain Cell Type Transcriptomes with Applications to Interpretation of Bulk Tissue Data. eneuro, pages ENEURO.0212–17.2017.

Mariani, J., Coppola, G., Zhang, P., Abyzov, A., Provini, L., Tomasini, L., Amenduni, M., Szekely, A., Palejev, D., Wilson, M., et al. (2015). Foxg1-dependent dysregulation of gaba/glutamate neuron differentiation in autism spectrum disorders. Cell, 162(2), 375–390.

McKenzie, M., Henders, A. K., Caracella, A., Wray, N. R., and Powell, J. E. (2014). Overlap of expression quantitative trait loci (eqtl) in human brain and blood. BMC medical genomics, 7(1), 31.

Newman, A.M., Liu, C.L., Green, M. R., Gentles, A. J., Feng, W., Xu, Y., Hoang, C. D., Diehn, M., and Alizadeh, A. A. (2015). Robust enumeration of cell subsets from tissue expression profiles. Nature Methods, 12(5), 453–457.

Olah, M., Patrick, E., Villani, A. C., Xu, J., White, C. C., Ryan, K. J., Piehowski, P., Kapasi, A., Nejad, P., Cimpean, M., Connor, S., Yung, C. J., Frangieh, M., McHenry, A., Elyaman, W., Petyuk, V., Schneider, J. A., Bennett, D. A., De Jager, P. L., and Bradshaw, E. M. (2018). A transcriptomic atlas of aged human microglia. Nat Commun, 9(1), 539.

Shabalin, A. A. (2012). Matrix eqtl: ultra fast eqtl analysis via large matrix operations. Bioinformatics, 28(10), 1353–1358.

Shen-Orr, S. S., Tibshirani, R., Khatri, P., Bodian, D. L., Staedtler, F., Perry, N. M., Hastie, T., Sarwal, M. M., Davis, M.M., and Butte, A.J. (2010). CelltypeâŁ”specific gene expression differences in complex tissues. Nature Methods, 7(4), 287–289.

Soreq, L., Rose, J., Soreq, E., Hardy, J., Trabzuni, D., Cookson, M. R., Smith, C., Ryten, M., Patani, R., Ule, J., *et al.* (2017). Major shifts in glial regional identity are a transcriptional hallmark of human brain aging. Cell reports, 18(2), 557–570.

Storey, J. D. and Tibshirani, R. (2003). Statistical significance for genomewide studies. Proceedings of the National Academy of Sciences, 100(16), 9440–9445.

Takasato, M., Pei, X. E., Chiu, H. S., Maier, B., Baillie, G. J., Ferguson, C., Parton, R. G., Wolvetang, E. J., Roost, M. S., de Sousa Lopes, S. M. C., et al. (2015). Kidney organoids from human ips cells contain multiple lineages and model human nephrogenesis. Nature, 526(7574), 564.

Wang, D., Liu, S., Warrell, J., Won, H., Shi, X., Navarro, F. C., Clarke, D., Gu, M., Emani, P., Yang, Y. T., et al. (2018). Comprehensive functional genomic resource and integrative model for the human brain. Science, 362(6420), eaat8464.

Wang, X., Park, J., Susztak, K., Zhang, N. R., and Li, M. (2019). Bulk tissue cell type deconvolution with multi-subject single-cell expression reference. Nat Commun, 10(1), 380.

Zeisel, A., Muñoz-Manchado, A. B., Codeluppi, S., Lönnerberg, P., La Manno, G., Juréus, A., Marques, S., Munguba, H., He, L., Betsholtz, C., et al. (2015). Cell types in the mouse cortex and hippocampus revealed by single-cell rna-seq. Science, 347(6226), 1138–1142.

