## Supplementary material for "Using multiple measurements of tissue to estimate subject- and cell-type-specific gene expression"

### Supplementary Figure



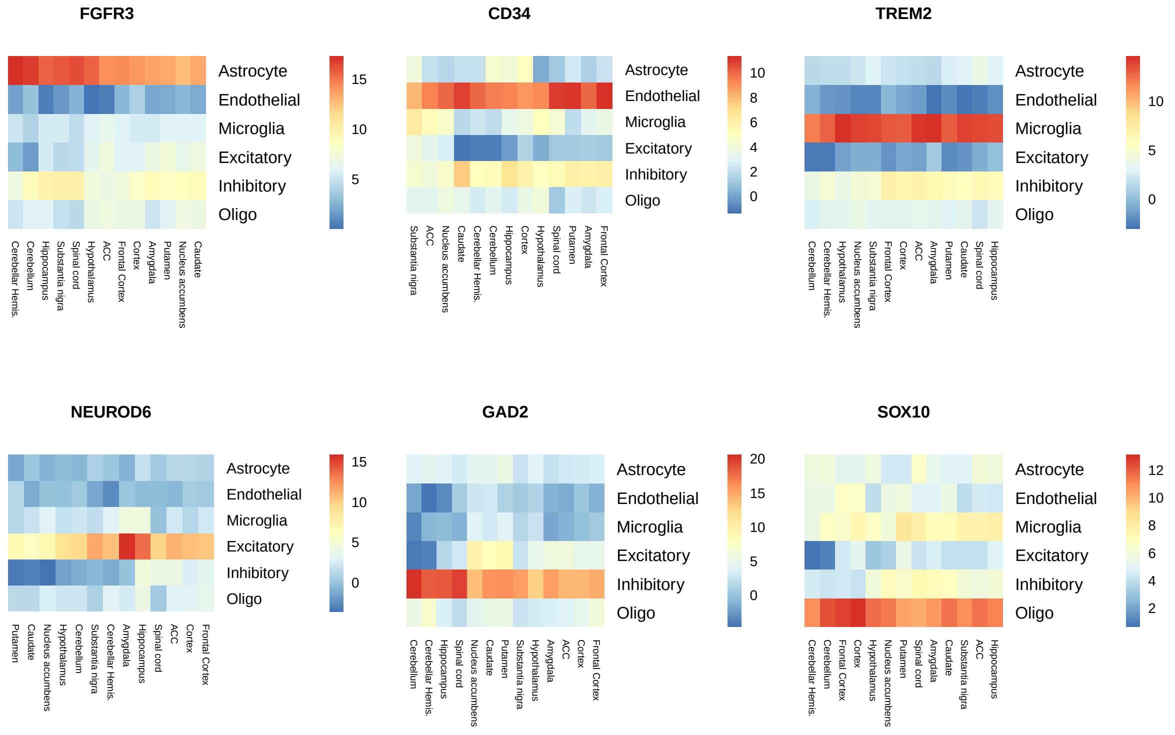


Supplementary Figure 1: Heatmaps of region-specific and CTS expression of marker genes estimated by reversing the role of subject and measure in MIND. The six marker genes correspond to astrocyte, endothelial cells, microglia, excitatory and inhibitory neurons, and oligodendrocyte (oligo), respectively.

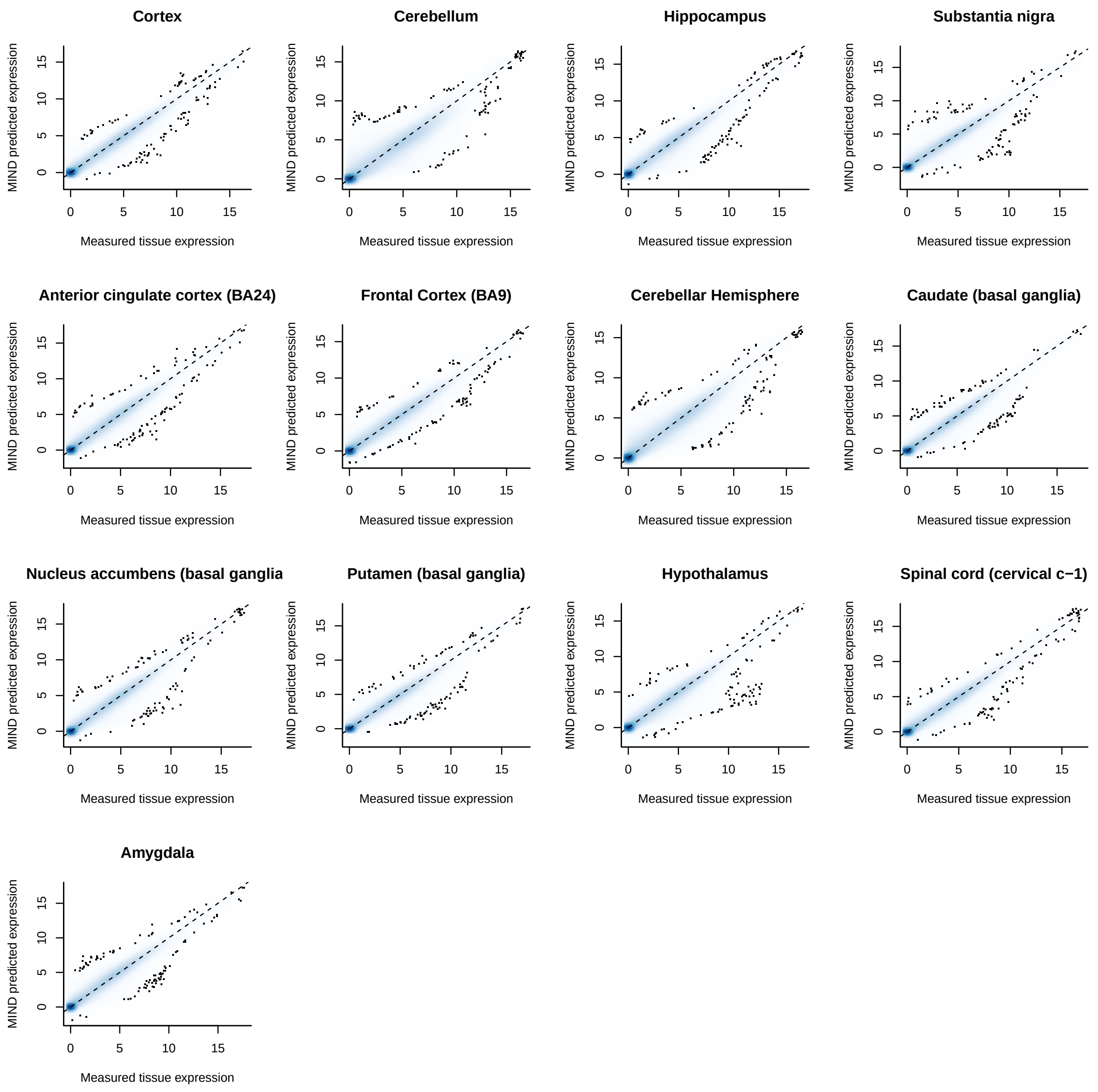


Supplementary Figure 2: Smoothed scatter plots of the observed GTEx brain tissue expression and MIND predicted expression for 13 GTEx brain regions. Dotted line at $y = x$. By the default setting of the smoothScatter function, the first 100 points from those areas of lowest regional densities are superimposed on the density image.

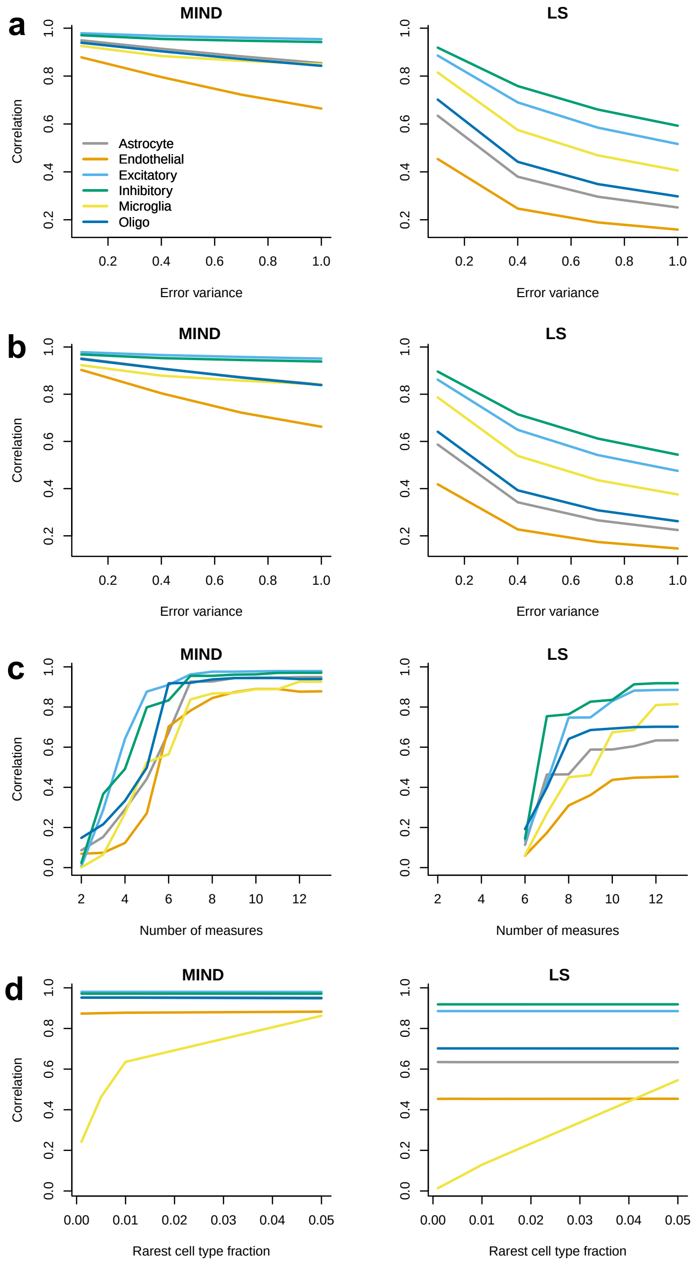


Supplementary Figure 3: Correlation between the true and MIND/LS (least squares) estimated expression for each of the six cell types in simulation. We simulated bulk expression data following equation (2) using the measured CTS expression (Habib *et al.*, 2017) and the estimated cell type fractions from GTEx brain data. We repeated the measured CTS expression from four subjects 25 times to simulate data for 100 subjects and then added independent measurement error to each. The results are based on 100 replications. (**a**) The impact of error variance. Here the number of measures is 13. (**b**) The impact of region-specific CTS expression. We added region-specific variation to CTS expression in addition to the simulation in (**a**). The variance of the region-specific variation is the same as the error variance. (**c**) The impact of the number of measures. Here the error variance is set as 0.1, which is what we observed in deconvolving GTEx brain data. LS is only available when the number of measures is equal to or more than the number of cell types. (**d**) The impact of rare cell type. We manipulated the fraction of microglia by making its mean value vary from 0.001 to 0.05, while keeping all other cell type fractions proportional to that observed in GTEx brain data.

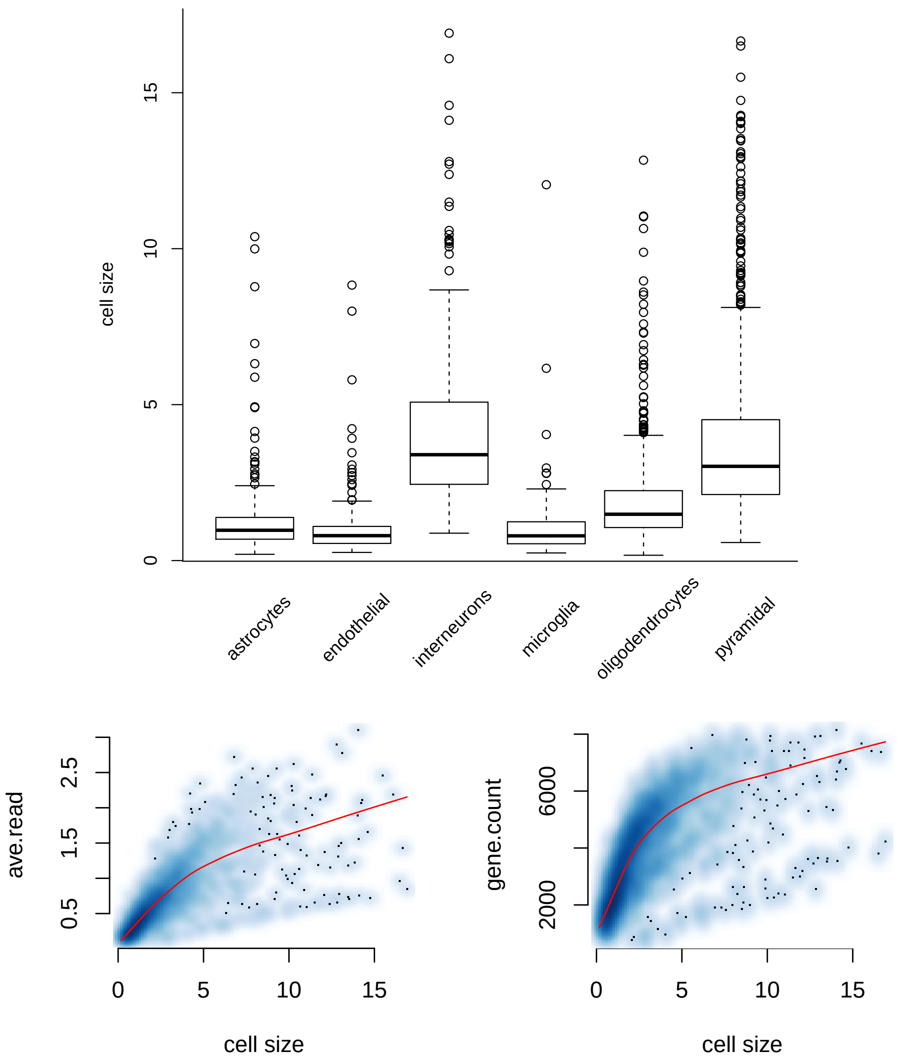


Supplementary Figure 4: The estimated cell size in the scRNA-seq data of Zeisel *et al.* (2015). Top: neurons (inhibitory/interneurons and excitatory/pyramidal neurons) have larger cell sizes as compared to non-neurons. Bottom: the average read (left) and gene count with nonzero read (right) vs. cell size. Both have a correlation of 0.6-0.7. The red line is a smooth curve.

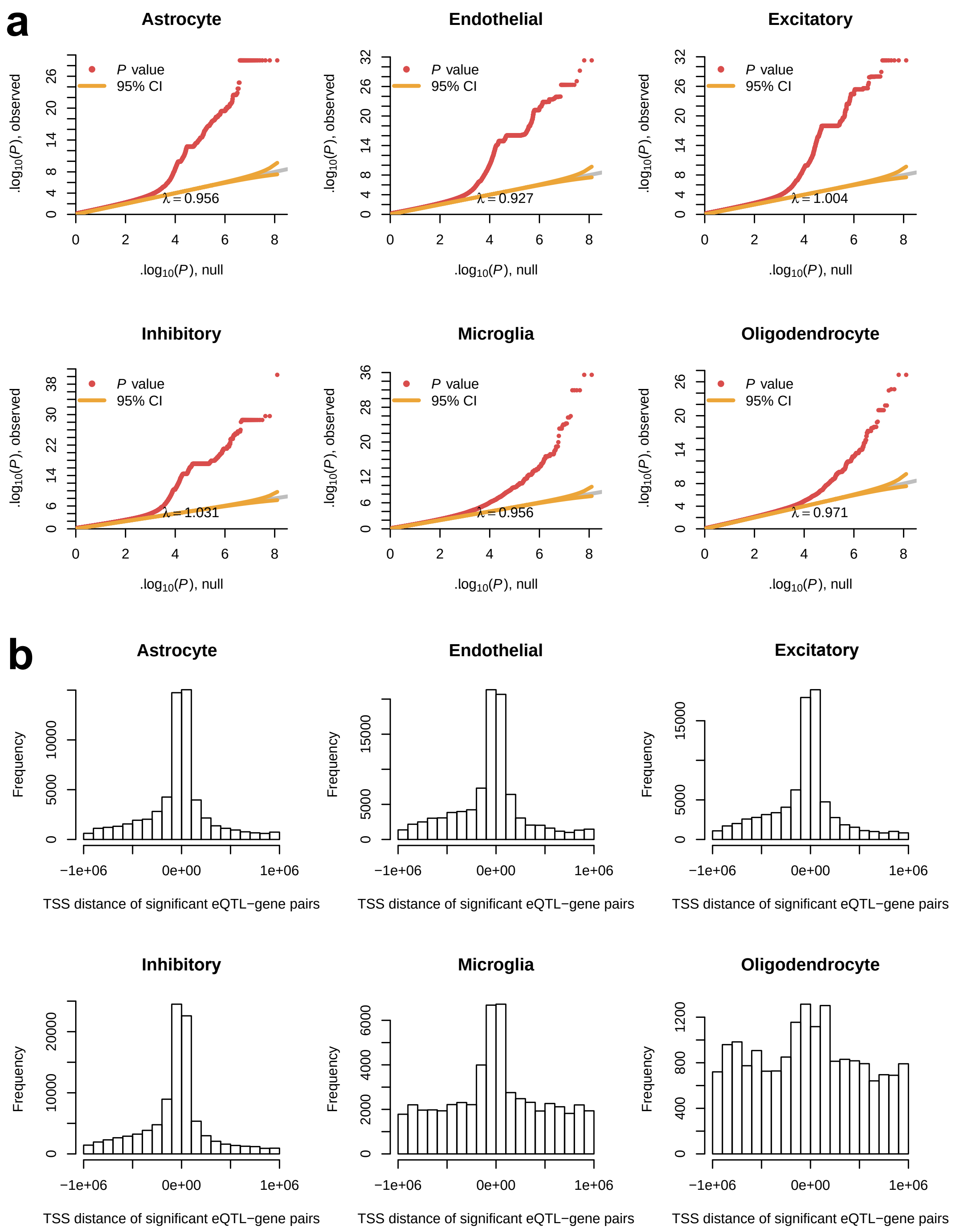


Supplementary Figure 5: Features of CTS eQTL mapping. (**a**) QQ-plots for eQTL mapping p-values for each cell type. (**b**) Enrichment of eQTLs near gene transcriptional start site (TSS) for each cell type. Note the concentration near the TSS except when the frequency of eQTLs discovered is relatively small for a cell type.

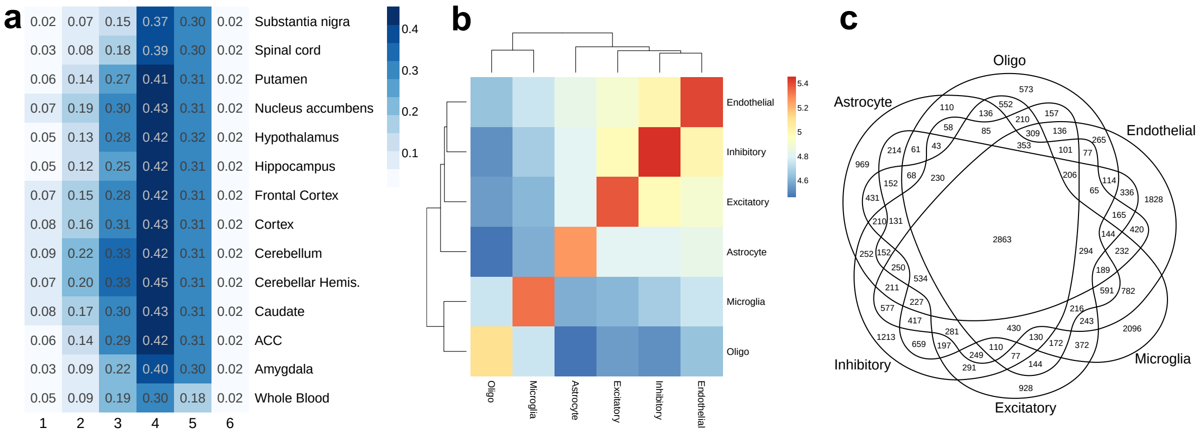


Supplementary Figure 6: (**a**) The probability of eQTLs identified in 1-6 cell types (columns) also being identified in each brain region or whole blood (rows). This is similar to Figure 5b but with printed probabilities. (**b**) The number of shared eQTLs for each pair of cell types (in log10 scale). (**c**) Venn diagram of eGenes (genes with eQTLs) identified in each cell type.

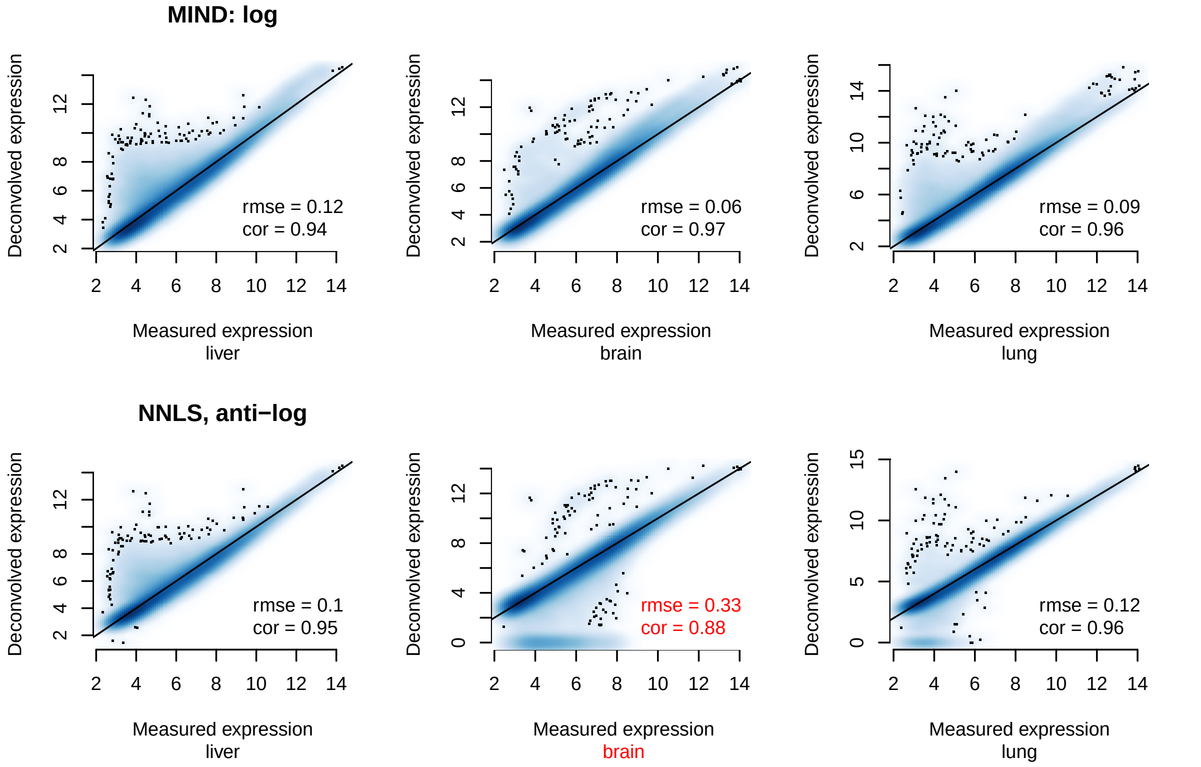


Supplementary Figure 7: Smoothed scatter plots comparing log and anti-log transformation in deconvolution using mixtures of tissue expression in Shen-Orr *et al.* (2010). There are three tissue types mixed: liver, brain, and lung. NNLS: non-negative least squares; rmse: root mean square error; cor: Pearson correlation.

### Supplementary Table

2

Supplementary Table 1: Computation time of MIND for varying numbers of subjects ($n$) and measures ($T$). Through simulations, we vary the number of subjects from 50 to 500, while keeping the number of measures as 10; we also vary the number of measures from 10 to 50, while keeping the number of subjects as 100. The memory usage is within 2GB. The testing is conducted using a single node on a laptop with Intel CORE i7 CPU. Note that MIND has built-in parallel computation across subjects to further speed up the computation.

| | T = 10 | | | | n = 100 | | |
|---|---|---|---|---|---|---|---|
| | n = 50 | 100 | 250 | 500 | T = 20 | 30 | 50 |
| time (min) | 1.6 | 3.2 | 8.2 | 19.0 | 4.7 | 6.2 | 7.9 |

Supplementary Table 2: Signature matrix estimated from Darmanis *et al.* (2015) in Excel file.

Supplementary Table 3: Average cell type fractions per region for GTEx brain data.

| | Astrocyte | Endothelial | Microglia | Excitatory | Inhibitory | Oligo |
|---|---|---|---|---|---|---|
| Putamen | 0.31 | 0.22 | 0.05 | 0.09 | 0.16 | 0.16 |
| Caudate | 0.32 | 0.21 | 0.07 | 0.11 | 0.19 | 0.10 |
| Amygdala | 0.25 | 0.18 | 0.07 | 0.17 | 0.20 | 0.14 |
| Nucleus accumbens | 0.30 | 0.17 | 0.05 | 0.16 | 0.25 | 0.06 |
| Hippocampus | 0.16 | 0.20 | 0.08 | 0.19 | 0.16 | 0.20 |
| Substantia nigra | 0.21 | 0.22 | 0.14 | 0.03 | 0.16 | 0.24 |
| Hypothalamus | 0.17 | 0.20 | 0.10 | 0.08 | 0.33 | 0.13 |
| Cortex | 0.22 | 0.18 | 0.03 | 0.34 | 0.20 | 0.03 |
| Frontal Cortex | 0.18 | 0.16 | 0.03 | 0.37 | 0.24 | 0.03 |
| ACC | 0.25 | 0.17 | 0.03 | 0.28 | 0.24 | 0.04 |
| Cerebellar Hemis. | 0.09 | 0.17 | 0.03 | 0.32 | 0.33 | 0.06 |
| Cerebellum | 0.15 | 0.18 | 0.03 | 0.30 | 0.29 | 0.05 |
| Spinal cord | 0.11 | 0.21 | 0.25 | 0.01 | 0.07 | 0.35 |

Supplementary Table 4: The correlation between measured and MIND estimated CTS expression in marker genes. For each cell type, we calculated the correlation between MIND estimated and Habib *et al.* (2017) measured expression. The maximum correlation in each column is in boldface. We can see that MIND has specificity to estimate the CTS expression for the targeted cell type.

| CTS expression in Habib *et al.* (2017) | MIND estimated CTS expression | | | | | |
|---|---|---|---|---|---|---|
| | Astrocyte | Endothelial | Excitatory | Inhibitory | Microglia | Oligo |
| Astrocyte | **0.65** | 0.24 | 0.54 | 0.38 | 0.00 | 0.48 |
| Endothelial | 0.27 | **0.57** | 0.31 | 0.19 | 0.08 | 0.27 |
| Excitatory | 0.38 | 0.24 | **0.75** | 0.56 | -0.15 | 0.49 |
| Inhibitory | 0.34 | 0.19 | 0.65 | **0.63** | -0.20 | 0.44 |
| Microglia | 0.03 | 0.02 | 0.04 | 0.06 | **0.49** | 0.16 |
| Oligo | 0.39 | 0.31 | 0.52 | 0.40 | 0.05 | **0.79** |

Supplementary Table 5: Analysis of simulated data mimicking bulk gene expression using MIND. To simulate bulk data, we use cell type fractions estimated in GTEx brain data. The profile matrix is built from Darmanis *et al.* (2015), either the signature matrix for marker genes or the profile matrix for all genes. For each simulation setting (each row), we vary the true value of variance parameters, $\sigma_e^2$, $\sigma_c^2$, and $\sigma_c^{kk'}$, which denote the error variance, and the variance and covariance of CTS expression, respectively. We present the average estimates of variance parameters and the correlation between the estimated (est.) and true CTS expression. The correlation is calculated for each cell type: astrocyte (astro), endothelial (endo), excitatory (excit) and inhibitory (inhit) neurons, microglial, and oligodendrocyte (oligo). The results are based on 100 replications.

| **Marker genes** | | | | | | | | | | | |
|---|---|---|---|---|---|---|---|---|---|---|---|
| true value | | | parameter estimate | | | correlation of est. and true CTS expression | | | | | |
| $\sigma_e^2$ | $\sigma_c^2$ | $\sigma_c^{kk'}$ | $\hat{\sigma}_e^2$ | $\hat{\sigma}_c^2$ | $\hat{\sigma}_c^{kk'}$ | astro | endo | excit | inhib | microglia | oligo |
| 1 | 1 | 0.5 | 1.00 | 1.05 | 0.48 | 0.97 | 0.96 | 0.98 | 0.97 | 0.94 | 0.97 |
| 2 | 2 | 1.0 | 1.99 | 1.93 | 0.99 | 0.95 | 0.93 | 0.97 | 0.95 | 0.90 | 0.94 |
| 3 | 3 | 1.5 | 3.00 | 2.69 | 1.52 | 0.93 | 0.90 | 0.95 | 0.93 | 0.86 | 0.92 |
| 4 | 4 | 2.0 | 4.00 | 3.41 | 2.05 | 0.92 | 0.87 | 0.94 | 0.92 | 0.83 | 0.90 |
| 5 | 5 | 2.5 | 5.00 | 4.12 | 2.59 | 0.90 | 0.85 | 0.93 | 0.90 | 0.80 | 0.88 |
| **All genes** | | | | | | | | | | | |
| true value | | | parameter estimate | | | correlation of est. and true CTS expression | | | | | |
| $\sigma_e^2$ | $\sigma_c^2$ | $\sigma_c^{kk'}$ | $\hat{\sigma}_e^2$ | $\hat{\sigma}_c^2$ | $\hat{\sigma}_c^{kk'}$ | astro | endo | excit | inhib | microglia | oligo |
| 1 | 1 | 0.5 | 1.00 | 1.04 | 0.48 | 0.97 | 0.95 | 0.97 | 0.96 | 0.95 | 0.96 |
| 2 | 2 | 1.0 | 1.99 | 1.93 | 0.99 | 0.94 | 0.90 | 0.94 | 0.93 | 0.91 | 0.93 |
| 3 | 3 | 1.5 | 3.00 | 2.67 | 1.52 | 0.92 | 0.87 | 0.92 | 0.91 | 0.88 | 0.90 |
| 4 | 4 | 2.0 | 4.00 | 3.38 | 2.06 | 0.90 | 0.84 | 0.91 | 0.89 | 0.85 | 0.88 |
| 5 | 5 | 2.5 | 5.00 | 4.09 | 2.59 | 0.88 | 0.81 | 0.89 | 0.87 | 0.82 | 0.86 |

### Supplementary Note

#### An EM algorithm for the multi-measure deconvolution model

Let $\boldsymbol{A}_{ij} = \boldsymbol{a}_j + \boldsymbol{B}_{ij}$, where $\boldsymbol{B}_{ij} \sim N(\boldsymbol{0}, \boldsymbol{\Sigma}_c)$. The complete-data log-likelihood is given by

$$\ell\left(\boldsymbol{a}, \Sigma_c, \sigma_e^2\right) = const - \frac{1}{2} np\log|\Sigma_c| - \frac{1}{2}\sum_{i=1}^{n}\sum_{j=1}^{p} \boldsymbol{B}_{ij}^{'} \Sigma_c^{-1} \boldsymbol{B}_{ij}$$

$$-\frac{p}{2}\sum_{i=1}^{n} T_i \log(\sigma_e^2) - \frac{1}{2\sigma_e^2}\sum_{i=1}^{n}\sum_{j=1}^{p}\left(\boldsymbol{X}_{ij} - \boldsymbol{W}_i\boldsymbol{a}_j - \boldsymbol{W}_i\boldsymbol{B}_{ij}\right)^{'}\left(\boldsymbol{X}_{ij} - \boldsymbol{W}_i\boldsymbol{a}_j - \boldsymbol{W}_i\boldsymbol{B}_{ij}\right).$$

##### E-step

The E-step is to calculate the expected value of the above statistics given the observed data and the current parameter estimates ($\boldsymbol{\gamma}^{(t)} = (\boldsymbol{a}^{(t)}, \Sigma_c^{(t)}, \sigma_e^{2(t)})$),

$$E\left(\ell\left(\boldsymbol{a}, \Sigma_c, \sigma_e^2\right) | \boldsymbol{X}, \boldsymbol{\gamma}^{(t)}\right) = const - \frac{p}{2}\sum_{i=1}^{n} T_i \log(\sigma_e^2) - \frac{1}{2} np\log|\Sigma_c|$$

$$-\frac{1}{2\sigma_e^2}\sum_{i=1}^{n}\sum_{j=1}^{p}\left[E\left(\boldsymbol{e}_{ij}^{'}|\boldsymbol{X}_{ij}, \boldsymbol{\gamma}^{(t)}\right) E\left(\boldsymbol{e}_{ij}|\boldsymbol{X}_{ij}, \boldsymbol{\gamma}^{(t)}\right) + \mathrm{tr}\left(\mathrm{var}\left(\boldsymbol{e}_{ij}|\boldsymbol{X}_{ij}, \gamma^{(t)}\right)\right)\right]$$

$$-\frac{1}{2}\sum_{i=1}^{n}\sum_{j=1}^{p}\left[\boldsymbol{B}_{ij}^{(t)'}\Sigma_c^{-1}\boldsymbol{B}_{ij}^{(t)} + \mathrm{tr}\left(\Sigma_c^{-1}\Sigma_{ij}^{(t)}\right)\right],$$

where

$$\boldsymbol{B}_{ij}^{(t)} = E\left(\boldsymbol{B}_{ij}|\boldsymbol{X}_{ij}, \boldsymbol{\gamma}^{(t)}\right) = \Sigma_c^{(t)}\boldsymbol{W}_i^{'}\left(\boldsymbol{W}_i\Sigma_c^{(t)}\boldsymbol{W}_i^{'} + \sigma_e^{2(t)} I_{T_i}\right)^{-1}\left(\boldsymbol{X}_{ij} - \boldsymbol{W}_i\boldsymbol{a}_j\right)$$
$$= \Sigma_{ij}^{(t)}\boldsymbol{W}_i^{'}\left(\boldsymbol{X}_{ij} - \boldsymbol{W}_i\boldsymbol{a}_j\right)/\sigma_e^{2(t)}$$

is the empirical Bayes estimate of $\boldsymbol{B}_{ij}$ and its covariance matrix is

$$\Sigma_{ij}^{(t)} = \mathrm{var}\left(\boldsymbol{B}_{ij}|\boldsymbol{X}_{ij}, \boldsymbol{\gamma}^{(t)}\right) = \Sigma_c^{(t)} - \Sigma_c^{(t)}\boldsymbol{W}_i^{'}\left(\boldsymbol{W}_i\Sigma_c^{(t)}\boldsymbol{W}_i^{'} + \sigma_e^{2(t)} I_{T_i}\right)^{-1}\boldsymbol{W}_i\Sigma_c^{(t)}$$
$$= \left(\boldsymbol{W}_i^{'}\boldsymbol{W}_i/\sigma_e^{2(t)} + \left(\Sigma_c^{(t)}\right)^{-1}\right)^{-1}.$$

For the error term,

$$E\left(\boldsymbol{e}_{ij}|\boldsymbol{X}_{ij}, \boldsymbol{\gamma}^{(t)}\right) = \sigma_e^{2(t)}\left(R_{ij}^{(t)}\right)^{-1}\left(\boldsymbol{X}_{ij} - \boldsymbol{W}_i\boldsymbol{a}_j\right),$$
$$\mathrm{var}\left(\boldsymbol{e}_{ij}|\boldsymbol{X}_{ij}, \boldsymbol{\gamma}^{(t)}\right) = \sigma_e^{2(t)} I_{T_i} - \sigma_e^{4(t)}\left(R_{ij}^{(t)}\right)^{-1},$$

where $R_{ij}^{(t)} = R_i^{(t)} = \boldsymbol{W}_i\Sigma_c^{(t)}\boldsymbol{W}_i^{'} + \sigma_e^{2(t)} I_{T_i}$.

##### M-step

In the M-step, we derive the estimate of the covariance matrix of random effects as

$$\Sigma_c^{(t+1)} = \frac{1}{np}\sum_{i=1}^{n}\sum_{j=1}^{p}\left[\boldsymbol{B}_{ij}^{(t)}\boldsymbol{B}_{ij}^{(t)'} + \Sigma_{ij}^{(t)}\right].$$

The error variance estimate is

$$\sigma_e^{2(t+1)} = \frac{1}{p\sum_{i=1}^{n} T_i}\sum_{i=1}^{n}\sum_{j=1}^{p}\left[E\left(\boldsymbol{e}_{ij}^{'}|\boldsymbol{X}_{ij},\boldsymbol{\gamma}^{(t)}\right)E\left(\boldsymbol{e}_{ij}|\boldsymbol{X}_{ij},\boldsymbol{\gamma}^{(t)}\right) + \mathrm{tr}\left(\mathrm{var}\left(\boldsymbol{e}_{ij}|\boldsymbol{X}_{ij},\gamma^{(t)}\right)\right)\right].$$

The average CTS expression for gene $j$ (part of profile matrix) is estimated as

$$\boldsymbol{a}_j^{(t+1)} = \left(\sum_{i=1}^{n}\boldsymbol{W}_i^{'}\left(R_i^{(t)}\right)^{-1}\boldsymbol{W}_i\right)^{-1}\sum_{i=1}^{n}\boldsymbol{W}_i^{'}\left(R_i^{(t)}\right)^{-1}\boldsymbol{X}_{ij}.$$

The final estimate for CTS expression in subject $i$ and gene $j$ is $\hat{\boldsymbol{A}}_{ij} = \hat{\boldsymbol{a}}_j + \hat{\boldsymbol{B}}_{ij}$.

#### Discussion on log vs. anti-log transformation

Zhong and Liu (2012) raised a concern about using log-transformed data in deconvolution. Shen-Orr *et al.* (2012) provided convincing argument about using log-transformation in their response. In addition, Shannon *et al.* (2014) showed more accurate results when using quantile normalized and log-transformed data to estimate cell type fractions.

Here we further address this issue using the same data as Zhong and Liu (2012), i.e., the mixtures of tissue expression in liver, brain, and lung by Shen-Orr *et al.* (2010). There are 33 mixtures of the three tissues with known mixing fractions. We compare the measured and deconvolved expression, for MIND using log-transformed data and NNLS (non-negative least squares) using anti-log transformed data (Supplementary Fig. 7). In MIND, the problem is formulated as 33 measures from a single subject, and NNLS treats it as 33 samples. The goal is to estimate the expression for each of the three tissues. The two approaches are comparable in liver and lung, in terms of root mean square error (rmse) and correlation, but anti-log transformed data produce much worse results in brain, which is the focus of our paper. The reason is that NNLS with anti-log transformed data fails to accurately deconvolve some genes and forces 6% of deconvolved expression exactly as zero.

#### Remark on cell fraction and cell size

While our estimates of the abundance of neurons, for example, match previous findings, such estimates can be inconsistent with those from neuroanatomical and other direct studies of cell representation (Azevedo *et al.*, 2009, Pelvig *et al.*, 2008). To better understand the estimated cell type fractions, we studied the relationship between cell size and gene expression using techniques in Jia *et al.* (2017) and results from Zeisel *et al.* (2015). We find that the estimated cell size is highly positively correlated with level of gene expression (Supplementary Fig. 4), and neurons tend to have a larger cell size than non-neurons, which agrees with previous findings (Wang *et al.*, 2018). Thus, while most deconvolution studies present their results in terms of estimated fractions of cell types, we believe these methods, including MIND, estimate the fraction of RNA molecules from each cell type instead.

#### Acknowledgements

The Genotype-Tissue Expression (GTEx) Project was supported by the Common Fund of the Office of the Director of the National Institutes of Health (commonfund.nih.gov/GTEx). Additional funds were provided by the NCI, NHGRI, NHLBI, NIDA, NIMH, and NINDS. Donors were enrolled at Biospecimen Source Sites funded by NCI\Leidos Biomedical Research, Inc. subcontracts to the National Disease Research Interchange (10XS170), Roswell Park Cancer Institute (10XS171), and Science Care, Inc. (X10S172). The Laboratory, Data Analysis, and Coordinating Center (LDACC) was funded through a contract (HHSN268201000029C) to The Broad Institute, Inc. Biorepository operations were funded through a Leidos Biomedical Research, Inc. subcontract to Van Andel Research Institute (10ST1035). Additional data repository and project management were provided by Leidos Biomedical Research, Inc.(HHSN261200800001E). The Brain Bank was supported supplements to University of Miami grant DA006227. Statistical Methods development grants were made to the University of Geneva (MH090941 & MH101814), the University of Chicago (MH090951,MH090937, MH101825, & MH101820), the University of North Carolina - Chapel Hill (MH090936), North Carolina State University (MH101819),Harvard University (MH090948), Stanford University (MH101782), Washington University (MH101810), and to the University of Pennsylvania (MH101822). The datasets used for the analyses described in this manuscript were obtained from dbGaP at http://www.ncbi.nlm.nih.gov/gap through dbGaP accession number phs000424.v7.p2.


#### References


Azevedo, F. A., Carvalho, L. R., Grinberg, L. T., Farfel, J. M., Ferretti, R. E., Leite, R. E., Filho, W. J., Lent, R., and Herculano-Houzel, S. (2009). Equal numbers of neuronal and nonneuronal cells make the human brain an isometrically scaled-up primate brain. *The Journal of Comparative Neurology*, **513**(5), 532–541.

Darmanis, S., Sloan, S. A., Zhang, Y., Enge, M., Caneda, C., Shuer, L. M., Hayden Gephart, M. G., Barres, B. A., and Quake, S. R. (2015). A survey of human brain transcriptome diversity at the single cell level. *Proceedings of the National Academy of Sciences*, **112**(23), 7285–7290.

Habib, N., Avraham-Davidi, I., Basu, A., Burks, T., Shekhar, K., Hofree, M., Choudhury, S. R., Aguet, F., Gelfand, E., Ardlie, K., *et al.* (2017). Massively parallel single-nucleus rna-seq with dronc-seq. *Nature Methods*, **14**(10), 955–958.

Jia, C., Hu, Y., Kelly, D., Kim, J., Li, M., and Zhang, N. R. (2017). Accounting for technical noise in differential expression analysis of single-cell rna sequencing data. *Nucleic acids research*, **45**(19), 10978–10988.

Pelvig, D., Pakkenberg, H., Stark, A., and Pakkenberg, B. (2008). Neocortical glial cell numbers in human brains. *Neurobiology of Aging*, **29**(11), 1754–1762.

Shannon, C. P., Balshaw, R., Ng, R. T., Wilson-McManus, J. E., Keown, P., McMaster, R., McManus, B. M., Landsberg, D., Isbel, N. M., Knoll, G., *et al.* (2014). Two-stage, in silico deconvolution of the lymphocyte compartment of the peripheral whole blood transcriptome in the context of acute kidney allograft rejection. *PloS one*, **9**(4), e95224.

Shen-Orr, S. S., Tibshirani, R., Khatri, P., Bodian, D. L., Staedtler, F., Perry, N. M., Hastie, T., Sarwal, M. M., Davis, M. M., and Butte, A. J. (2010). Cell type–specific gene expression differences in complex tissues. *Nature Methods*, **7**(4), 287–289.

Shen-Orr, S. S., Tibshirani, R., and Butte, A. J. (2012). Gene expression deconvolution in linear space. *Nature methods*, **9**(1), 9.

Wang, J., Huang, M., Torre, E., Dueck, H., Shaffer, S., Murray, J., Raj, A., Li, M., and Zhang, N. R. (2018). Gene expression distribution deconvolution in single-cell rna sequencing. *Proceedings of the National Academy of Sciences*, **115**(28), E6437–E6446.

Zeisel, A., Muñoz-Manchado, A. B., Codeluppi, S., Lönnerberg, P., La Manno, G., Juréus, A., Marques, S., Munguba, H., He, L., Betsholtz, C., *et al.* (2015). Cell types in the mouse cortex and hippocampus revealed by single-cell rna-seq. *Science*, **347**(6226), 1138–1142.

Zhong, Y. and Liu, Z. (2012). Gene expression deconvolution in linear space. *Nature methods*, **9**(1), 8.
